# *In vivo* changes in zebrafish anesthetic sensitivity in response to the loss of *kif5Aa* are associated with the alteration of mitochondrial motility

**DOI:** 10.1101/2024.12.20.629838

**Authors:** Priya Dubey, Roshni Datta, Roderic G Eckenhoff, Victoria M Bedell

## Abstract

Anesthetic and sedative drugs are small compounds known to bind to hundreds of proteins. One intriguing binding partner of propofol is the kinesin motor domain, kif5A, a neuronal mitochondrial transport protein. Here, we used zebrafish WT and *kif5Aa* KO larval behavioral assays to assess anesthetic sensitivity and combined that with zebrafish primary neuronal cell culture to probe for alteration in mitochondrial motility. We found that the loss of *kif5Aa* increases behavioral sensitivity to propofol and etomidate, with etomidate hypersensitivity greater than propofol. In contrast, *kif5Aa* KO animals were resistant to the behavioral effects of dexmedetomidine. Finally, WT and *kif5Aa* KO larvae responded similarly to the behavioral effects of ketamine. Propofol inhibited the anterograde motility of mitochondria in WT zebrafish neurons, while etomidate inhibited mitochondrial motility in both anterograde and retrograde directions; neither drug altered mitochondrial motility in the *kif5Aa* knockout (KO) neurons. In contrast, dexmedetomidine enhanced retrograde mitochondrial motility in both WT and *kif5Aa* KO animals. Finally, ketamine had little significant effect on mitochondrial motility in either mutant or WT animals. These data demonstrate that each anesthetic/sedative drug affects the motor protein machinery uniquely and is associated with unique changes in behavior. Understanding how different anesthetic compounds alter neuron motor proteins will be important in defining how anesthetics alter neuronal signaling and energetic dynamics.

## Introduction

There is a remarkable variety of small compounds capable of producing changes in consciousness, ranging from sedation to general anesthesia. These drugs have many incompletely understood effects on the central nervous system (CNS), including energy production and use[1, 2]. The neuronal synapse has one of the highest energy requirements in the brain[3, 4] and, the highest mitochondrial concentration[5]. Additionally, mitochondria contribute to volatile anesthetic action[6] by specifically inhibiting complex I of the electron transport chain and decreasing synaptic ATP[7]. Decreased synaptic excitatory transmission is observed in animal models with anesthetic hypersensitivity caused by cell-specific impaired mitochondrial function[6, 7]. However, action upon the electron transport chain may not fully define anesthetics’ effect on cellular energy availability.

Complex systems have evolved to transport neuronal mitochondria from their site of biogenesis in the cell body[8] to and from the synapse[9], presumably to align with synaptic energy needs[10]. Kinesin (kif) is the motor protein family responsible for the anterograde movement of various cargoes within cells[11]. While over 80% of the kinesin superfamily are expressed in neurons[11], three kinesin families, Kif5, Kif1, Kinesin like protein 6 (Klp6) have been shown to transport mitochondria within the axons[12–14].

Of these kinesins, previous studies on propofol-binding partners identified kif5B as a target[15]. Propofol binds with high affinity to the motor head domain of kif5B, causing premature cargo detachment from the microtubule [15–17]. Additionally, the propofol binding pocket within kif5B has been identified as requiring two specific amino acids within the motor domain (**Supplemental Fig 1**). While the motor domain is largely conserved, it is still unknown if the IV anesthetics bind to other kinesins. Additionally, it remains unknown if changing kinesin motility alters behavioral outcomes *in vivo*. Therefore, to answer these questions, we employed the zebrafish model organism.

With the zebrafish, kif5 family expression patterns are known[18]. *kif5Ba* and *kif5Bb* are ubiquitously expressed within the developing animal. However, *kif5Aa* is primarily expressed within the nervous system and is a key mitochondrial transport protein[18–20]. Further, we found significant homology within the motor domain and propofol binding pocket between Kif5Aa and Kif5B (**Supplemental Fig 1**). Therefore, we decided to employ the *kif5Aa* knockout (KO) zebrafish line as an intersection between motor domain homology and mitochondrial motility.

The question of how the loss of *kif5Aa* affects larval movement and CNS mitochondrial motility is still an open one. We use whole animal behavioral analysis to assess whether sensitivity to four distinct intravenous (IV) anesthetic/sedative drugs is altered by the KO, and primary neuronal cell culture to assess for associated changes in mitochondrial motility.

## Methods

### Zebrafish husbandry

All zebrafish experiments complied with the University of Pennsylvania Institutional Animal Care and Use Committee (IACUC). The adult zebrafish were maintained in an aquatic facility overseen by the University Laboratory Animal Resources (ULAR) at the University of Pennsylvania.

All zebrafish were maintained at 28 °C with a 13-11 light-dark cycle as previously described[21]. Adult fish were maintained using standard husbandry conditions[21]. KO and WT siblings were maintained in the same conditions until experimentation.

### Genotyping the kif5Aa KO line

Genomic DNA was obtained from adult animals using the fin-clipping method[21]. The PCR identifying the kif5Aa^sa7168^ KO animals was run using previously described primers and digested using Dra III[19].

*kif5Aa^sa7168^* heterozygous zebrafish were mated, and before behavioural testing, the larvae were separated based on known visual phenotypes: loss of the swim bladder and increased pigmentation. “WT” were a combination of WT and heterozygous animals as they were indistinguishable.

The *kif5Aa* heterozygous line was outcrossed into a pan-mitochondrial Grx ro-GFP2 fluorescent line for mitochondrial tracking[22].

### Drug exposures

We tested four sedative/anesthetics: propofol, etomidate, dexmedetomidine, and ketamine. Each drug was diluted into E3 embryo water for the behavior experiments described previously[23]. Briefly, propofol was diluted to 100 μM from a 100 mM DMSO stock in E3 by sonicating 5 minutes (min); absorbance at 270nm was used to confirm the concentration. Etomidate and dexmedetomidine were solubilized directly by vortexing. Finally, ketamine was diluted at in E3 with 5mM hepes at a pH of 8, which is required for improved uptake by the larvae.

For cell culture, all drugs were initially diluted to a 2x concentration using 1x PBS. This was then diluted to the final drug concentration in complete cell culture media to ensure adequate nutrition for the cells.

### Zebrafish 5 dpf behavioural assay

Behavior experiments were performed as previously described[23]. Briefly, we used the DanioVision behavioral system with EthoVision software (Noldus) to measure the distance the WT and *kif5Aa* KO larvae moved spontaneously (SM) over 4 min or in 1 second after an elicited acoustic stimulus (EM). The experiments were 2 hours within the dark enclosure of the behaviour system, with the EM at the highest setting every 30 minutes. All experiments used 96-well glass plates (Chemglass Life Sciences, CG-1910) at 28°C.

### Primary zebrafish neuronal cell culture

This protocol is based on a previously described zebrafish neuronal cell culture assay[24]. Briefly, we used a 96-well tissue culture plate with a glass-like polymer coverslip bottom (Cellvis, P96-1.5P) coated with Entactin-Collagen-Laminin (EMD Millipore, 08-110) at 10-20 μg/mL and stored at 4 °C overnight.

Twenty 5 dpf zebrafish WT siblings and *kif5Aa* KO larvae containing the Grx ro-GFP2 were euthanized on ice until the heart stopped beating. Brains were dissected and placed in 500 μL Hanks base salt solution with 1% penicillin/streptomycin on ice. 200 μL of Accutase (Sigma, A6964) was added for dissociation and incubated for 10 min in a 37 °C water bath. Brains were triturated using a glass pipette for 2-4 min or until there were few visible clumps of cells. This was key to decreasing aggregation and promoting attachment and axon outgrowth during cell culturing. The dissociation reaction was stopped using 700 μL of complete media. Complete media consisted of 0.22 μm sterile-filtered Leibovitz-15, 2.5mM Glutamax I, 15 ng/mL epidermal growth factor, 10% fetal bovine serum, 1% penicillin/streptomycin and 5% zebrafish embryo extract. The embryo extract was created as previously described[25].

The sample was centrifuged at 1500 RPM for 6 min at 4 °C. The supernatant was discarded and 100-125 μL of complete media was added to resuspend the pellet. The cell mixture was plated using 50-75 μL per well. The cells were incubated at 28 °C for 2 days.

### Imaging

Neurons were imaged using a Leica DMI4000 with a Yokagawa CSU-X1 spinning disk confocal attachment at 63x.

The anaesthetic concentration chosen was the EC50s previously determined[23]: propofol 3 μM, etomidate 30 μM, dexmedetomidine 10 nM, and ketamine 130 μM. The cells were incubated in the drug for at least 30 min before imaging. Neurons that could be traced back to a specific cell body were imaged every 3 seconds for 5 minutes for a total of 101 images (initial image plus imaging for the next five minutes). Five to ten cells were imaged per well.

An open-source ImageJ plugin, KymoToolBox, defined mitochondrial motility. **Supplemental Table 1** is a summary of the mitochondria traced. Briefly, each movie contained a range of 1-6 axons. The axons were highlighted, and a kymograph was created, on which individual mitochondria were traced. The number of traceable mitochondria varied between movies, therefore, the range of mitochondria traced is given in **Supplemental Table 1**. Any movie with less than 5 traceable mitochondria was discarded. ImageJ was then able to calculate the distance moved for the mitochondria. Total mitochondrial movement is herein defined as any movement, retrograde plus anterograde, including zero movement. No minimum motility down the axon was used.

### Statistics

For behavior studies, the distance moved for 8-12 larvae per drug concentration per genotype was averaged and counted as a single experimental value (n). The experiment was performed at least 4 times on separate days to ensure technical and biological replication. Each animal was exposed to a single drug and not reused. For the baseline movement assays, 40-41 experiments at 30 minutes were analyzed and significance tested using the Mann-Whitney test for nonparametric data. Hill curves were created using 6-8 drug concentrations, including a no-drug (ND) control. The distance moved was normalized using the ND as 100% movement. Hill curves were created using GraphPad Prism. As with prior studies, the Hill curves were constrained to a top value of 100% and a bottom of 0%. The EC50s with 95% confidence intervals (CI) were calculated and plotted. EC50 comparison p values were determined using a sum of squares F test.

All experiments were performed at least in triplicate for the mitochondrial motility assays. The motility data was, in most cases, normally distributed. Therefore, a two-way ANOVA using Bonferroni’s correction for multiple comparisons determined significance. For all groups, significance is labeled: not significant (n.s., p > 0.05), p ≤ 0.05 (*), p ≤ 0.01 (**), p ≤ 0.001 (***), p ≤ 0.0001.

## Results

### Loss of *kif5Aa* alters behavioral sensitivity in a drug-specific manner

We examined larval movement using two previously described behavioral endpoints: loss of spontaneous movement (SM) and loss of elicited movement (EM) to an acoustic tap stimulus[23]. Initially, we tested baseline movement to the two behaviors at 5 dpf. The *kif5Aa* KO larvae significantly increased SM compared to their wild-type siblings (WT) (**Supplemental Fig. 2A**). In contrast, the *kif5Aa* KO animals showed significantly decreased EM (**Supplemental Fig. 2B**). For all subsequent studies, movement of no drug controls was defined as 100%.

Because it was unclear if the depletion of synaptic mitochondria was a time-dependent phenomenon during the maintenance phase of anesthesia, we decided to test for changes in sensitivity over time. First, we assessed *in vivo* sensitivity to propofol over time in the 5 dpf larvae. Using SM, the *kif5Aa* KO was more sensitive to propofol than their WT siblings (**Fig. 1A-D)**), with a 30-40% decrease in the EC50 (**Fig. 1E**). Using EM, there was no initial difference in sensitivity (**Fig. 1F-J**), but after 90 minutes of exposure, the *kif5Aa* KO became significantly more sensitive to propofol than WT (**Fig. 1H-I**). We were able to see a decrease in the EC50 of the *kif5Aa* KO larvae over time (**Fig. 1J**).

**Figure 1.**
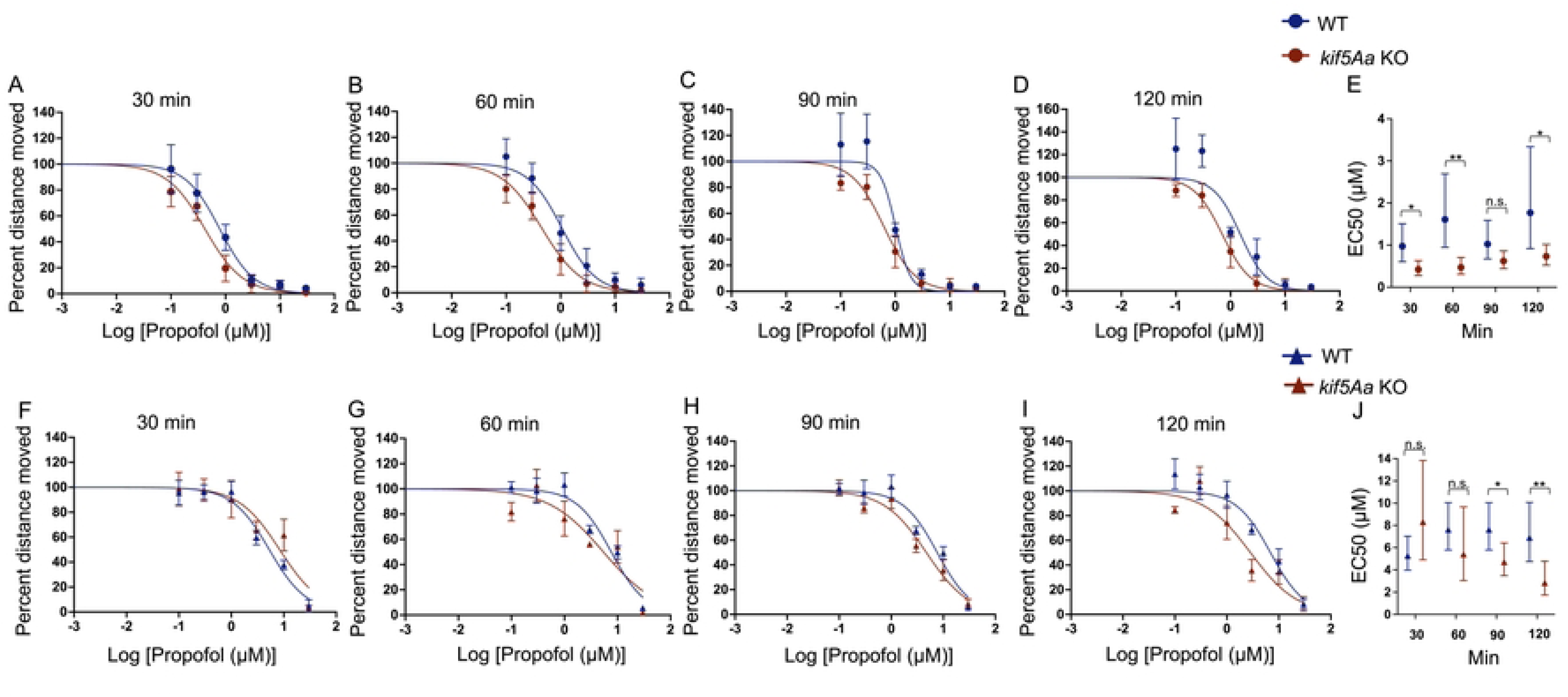
*kif5Aa* KO 5dpf larvae are more sensitive to propofol. A-D) Increased sensitivity to propofol in the *kif5Aa* KO larvae (red) as compared to WT (blue) using spontaneous movement (SM) is consistent over time. E) A significant decrease in the EC50 of *kif5Aa* KO versus WT over time. F-G) No significant change in sensitivity was seen with elicited movement. H-I) By 90 minutes, there was a significant sensitivity to propofol with EM. J) The EC50s within *kif5Aa* KO larvae decreased over time and showed sensitivity to propofol at 90- and 120-min. Hill curves: 7 propofol concentrations (0-30 mM), n = 7-9 (average of 8-12 zebrafish per genotype per drug dose), SEM error bars. EC50s: calculated from Hill curves, statistical significance using the extra sum of squares F test, 95% CI error bars.

Next, we assessed sensitivity to etomidate and found *kif5Aa* KO larvae were 85-90% more sensitive using the SM endpoint compared to the WT siblings (**Fig. 2A-E**). When using the EM endpoint, like propofol, when exposed to etomidate the *kif5Aa* KO larvae had no significant sensitivity at 30-60 minutes (**Fig. 2F-G**). However, by 90 minutes *kif5Aa* KO larvae were significantly more sensitive to etomidate (**Fig. 2H-I**). The larvae demonstrated a slow, time-dependent increase in sensitivity to etomidate (**Fig 2J**).

**Figure 2.**
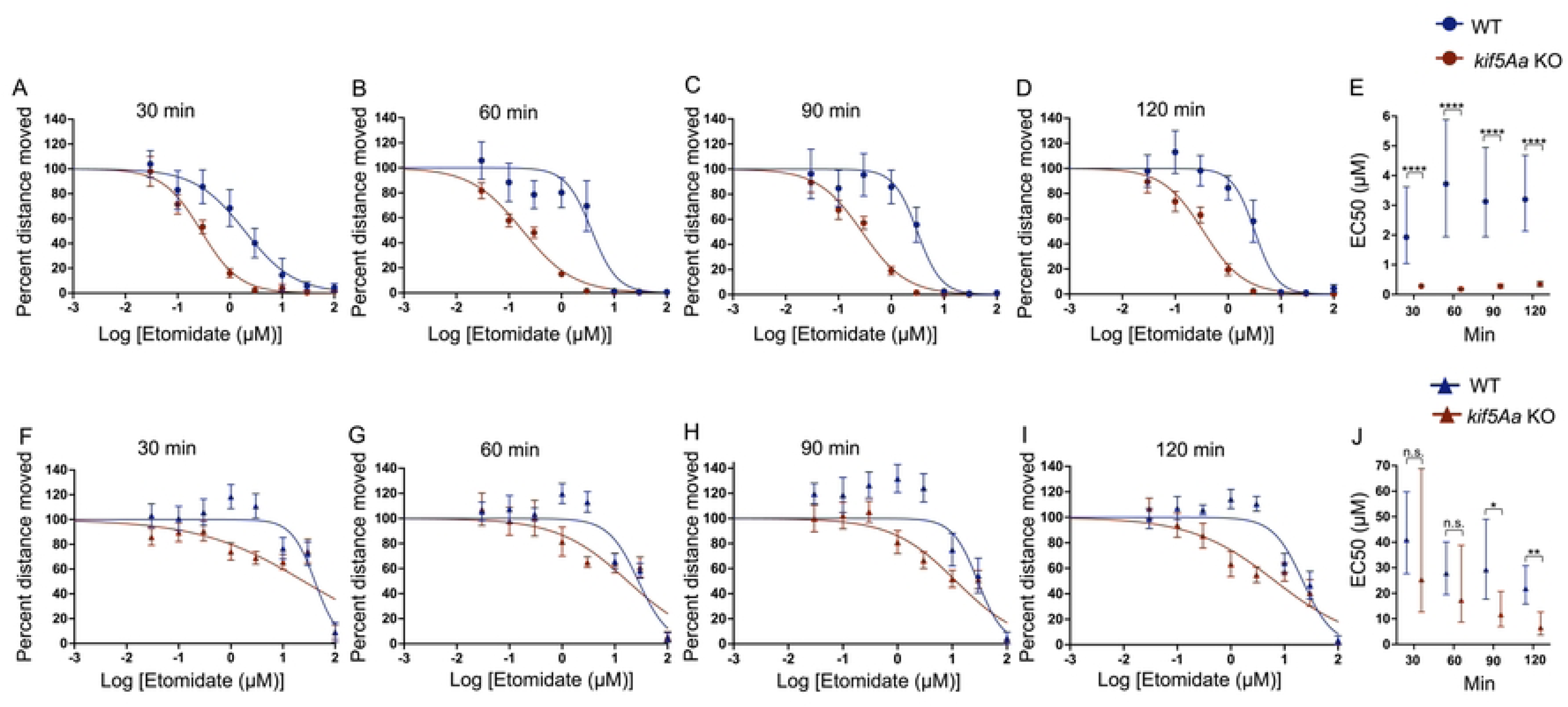
*kif5Aa* KO 5 dpf larvae are more sensitive to etomidate. A-D) When testing spontaneous movement (SM), the *kif5Aa* KO larvae (red) were more sensitive to etomidate as compared to their WT siblings (blue). This sensitivity was consistent over time. E) There was a significant decrease in the EC50 of *kif5Aa* KO compared to WT siblings, which is consistent over time. F-G) Elicited movement (EM) showed no significant difference in *Kif5Aa* KO compared to WT. H-I) At 90 and 120 minutes, there were significant increase in sensitivity in *kif5Aa* KO larvae vs WT siblings. J) Time-dependent decrease in EC50s in *kif5Aa* KO larvae, which became significant at 90 and 120 minutes. Hill curves: 9 etomidate concentrations (0 to 100 mM), n = 7-13 (average of 8-12 larval zebrafish per n), SEM. EC50s: calculated from the Hill curves, statistical significance using the extra sum of squares F test, 95% CI error bars.

Given the consistent SM behavioral sensitivity, we examined a sedative drug without full anesthetic properties, dexmedetomidine. We found a time-dependent resistance to dexmedetomidine in the kif5Aa larvae (**Fig. 3A-C**) that resolved by 120 minutes (**Fig. 3D**). We found a 2-4-fold increase in the EC50 for the first 90 minutes (**Fig. 3E**). Consistent with dexmedetomidine being only sedative, there was no loss of movement to the EM (**Fig. 3F-I**).

**Figure 3.**
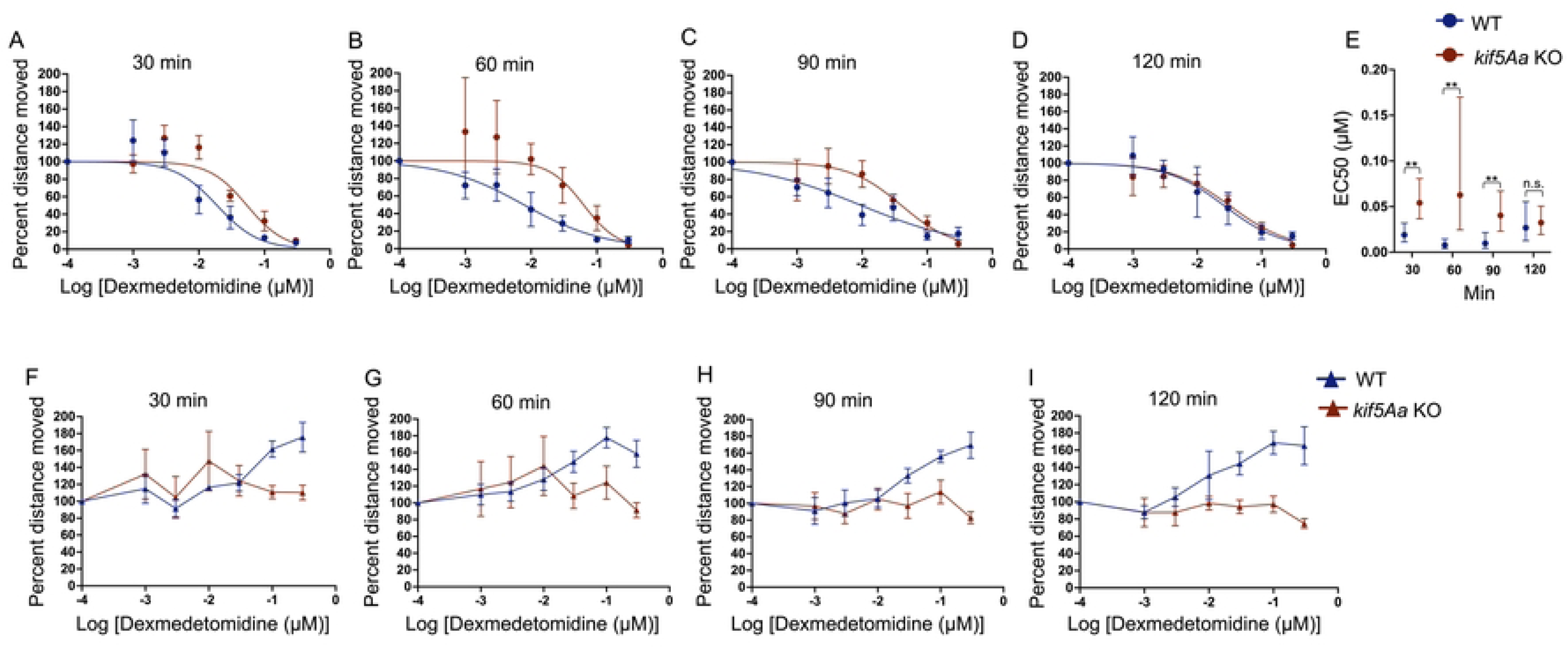
*kif5Aa* KO 5 dpf larvae are less sensitive to dexmedetomidine. A-D) When testing spontaneous movement (SM), kif5Aa KO larvae (red) were found to be resistant to dexmedetomidine as compared to their WT siblings (blue). D) This resistance resolved over time. E) The EC50s showed a significant increase in the *kif5Aa* KO larvae as compared to WT, which resolved by 120 min. H-I) Dexmedetomidine was a known sedative that did not induce general anesthesia. There was no loss of movement to the EM. At supratherapeutic doses of dexmedetomidine (0.1 and 0.3 mM, last two points on the blue line), there was an increase in WT movement that was not seen in the *kif5Aa* KO larvae. Hill curves: 7 dexmedetomidine concentrations (0 to 0.3 mM), SEM error bars. EC50s: n = 5-8, (average of 8-12 larval zebrafish per genotype per drug dose). EC50s: calculated from the Hill curves, statistical significance using the extra sum of squares F test, 95% CI error bars.

Ketamine, a non-GABAergic anaesthetic, produced no change in the loss of SM (**Fig. 4A-D**) and the EC50s of WT to *kif5Aa* KO larvae overlapped (**Fig. 4E**). Similarly, there was no change in response to EM when exposed to ketamine (**Fig. 4F-J**).

**Figure 4.**
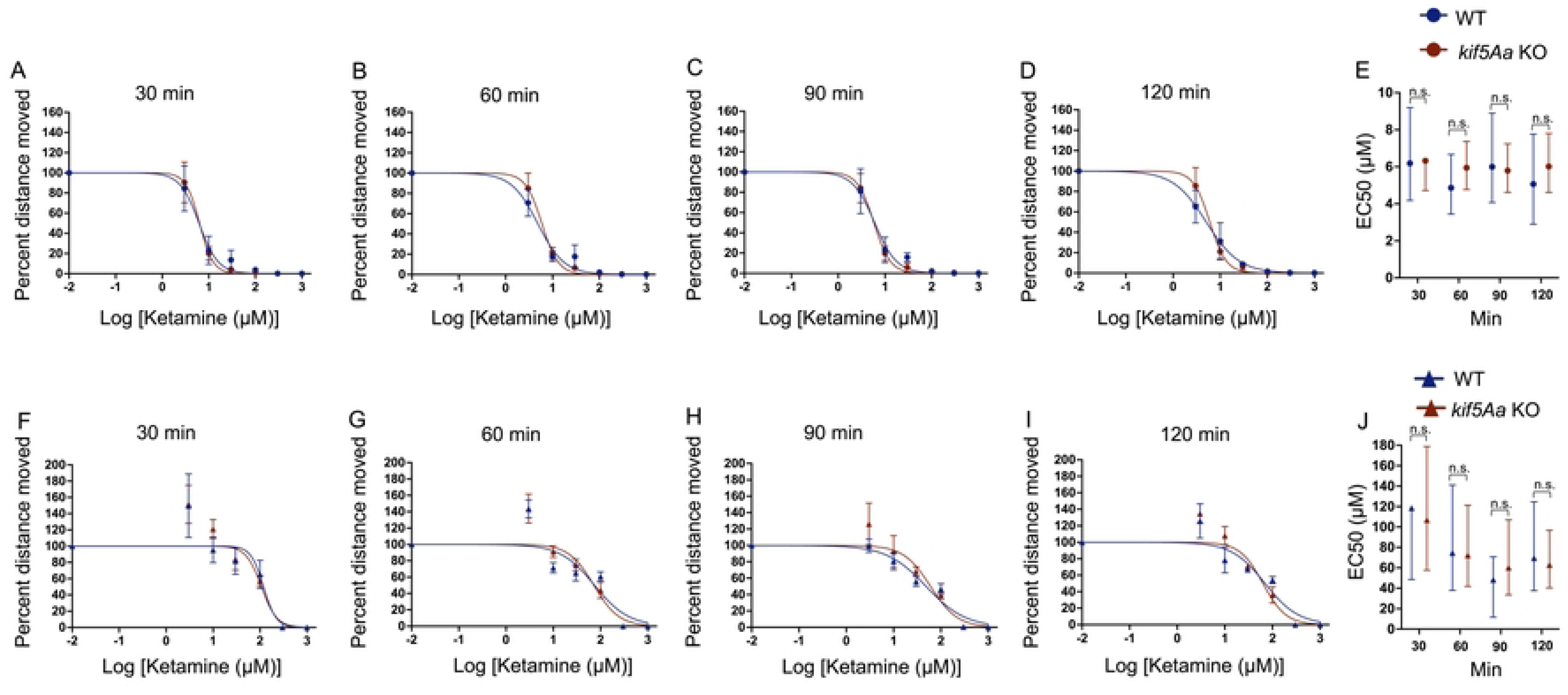
Ketamine does not change behavioral sensitivity in 5 dpf *kif5Aa* KO larvae. A-D) When testing SM, the Hill curve does not change in response to ketamine. E) The EC50s of WT (blue) and *kif5Aa* KO (red) overlap. F-I) Similarly, there was no change in EM in response to ketamine. J) The EC50s of *kif5Aa* KO (red) and their WT siblings (blue) are equivalent. Due to the lack of phenotype, it was decided to preserve animals and not further replicate the data to improve the 95% CI of the EC50s.

### Mitochondrial motility is altered in a drug-specific manner

We next asked whether alterations in anesthetic sensitivity were associated with changes in mitochondrial motility within primary neuron cells cultured from both WT (**Supplemental Fig. 3A**) and *kif5Aa* KO (**Supplemental Fig. 3B**) brains. All mitochondria within the neuron were labeled, showing the interconnected network the organelle creates. Time-lapse moves were created of the neurons, and the axons were highlighted in Image J (**Supplemental Fig. 4A-A’; B-B’;** WT **Supplemental Movie1**, *kif5Aa* KO **Supplemental Movie2)**. Using these movies, kymographs were created, and the mitochondria were traced over time (**Supplemental Fig. 4A’’; B’’**).

Baseline mitochondrial motility with no drug (ND) was significantly different in the *kif5Aa* KO neurons as compared to the WT for every measurement, including motility, distance moved, and velocity. Further, this difference was seen in total movement, anterograde and retrograde (first ND blue bar and first ND red bar, **Figs. 5-8**).

**Figure 5.**
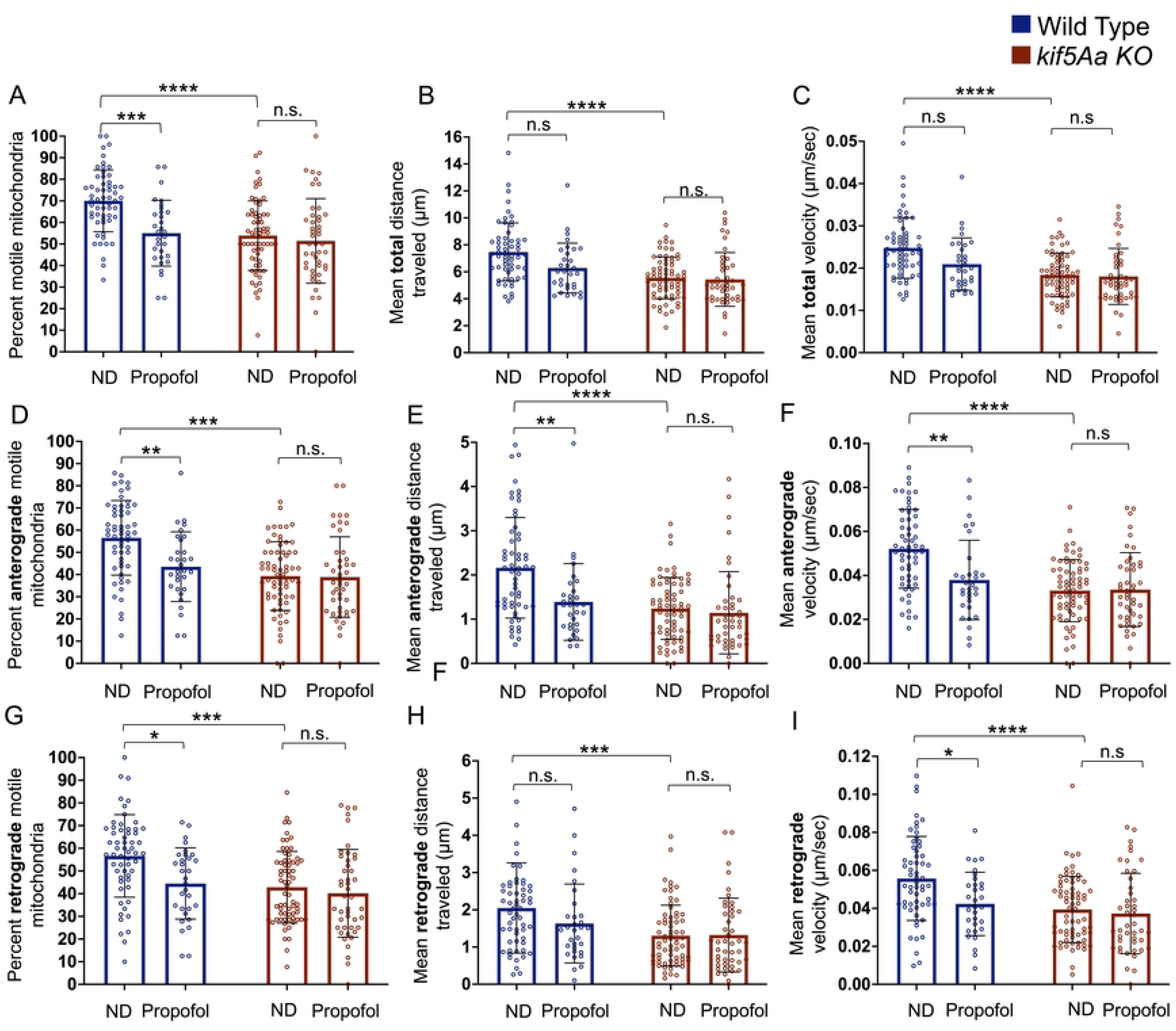
Exposure to propofol primarily decreases anterograde mitochondrial motility in WT neurons. Mitochondrial motility was assessed in cultured primary zebrafish neurons. When comparing no drug (ND) WT to *kif5Aa* KO mitochondrial movement, there was a decrease in the percent of motile mitochondrial, distance moved, and velocity despite directionality (A-I, first red bar and first blue bar). A) There was a decrease in the percent of motile mitochondria in WT neurons when exposed to propofol (blue bars). B-C) There was no change in the total distance moved or velocity (blue bars) within WT neurons. D-F) When assessing anterograde motility in WT neurons, there was a significant decrease in the number of motile mitochondria, the distance moved, and velocity (blue bars). G) In WT neurons, there was a decrease in the number of motile retrograde mitochondria (blue bars), (H) no change in the distance moved (blue bars) but a (I) decrease in retrograde velocity (blue bars). A-I) When *kif5Aa* KO neurons were exposed to propofol, there was no significant change in movement (red bars). n= 31-68 (see Supplemental Table 2). Error was in standard deviation and statistical significance using a two-way ANOVA with multiple comparisons using Bonferroni’s correction.

In WT neurons, propofol produced a significant decrease in the total number of motile mitochondria (**Fig. 5A**, blue bars) without a change in the total distance moved (**Fig. 5B**, blue bars) or in the total velocity (**Fig. 5C**, blue bars). Dividing the total movement by direction, propofol produced a significant decrease in anterograde mitochondrial motility, distance moved, and velocity (**Fig. 5D-F**, blue bars). There was also a decrease in retrograde motility and velocity (**Fig. 5G**, **4I**, blue bars) but no change in retrograde distance (**Fig. 5H**, blue bars). In contrast, propofol produced no motility changes in *kif5Aa* KO neurons (**Fig. 5A-I**, red bars).

Etomidate significantly reduced the number of motile mitochondria (**Fig. 6A**, blue bars), the total distance mitochondria moved (**Fig. 6B**, blue bars), and the mitochondrial velocity (**Fig. 6C**, blue bars) in WT neurons. Dividing into anterograde and retrograde directions, etomidate produced a significant decrease in the number of motile mitochondria, distance moved, and velocity in both directions (**Fig. 6E-I**, blue bars). Etomidate produced no change in the *kif5Aa* KO neuron mitochondrial motility (**Fig. 6A-I, red bars**).

**Figure 6.**
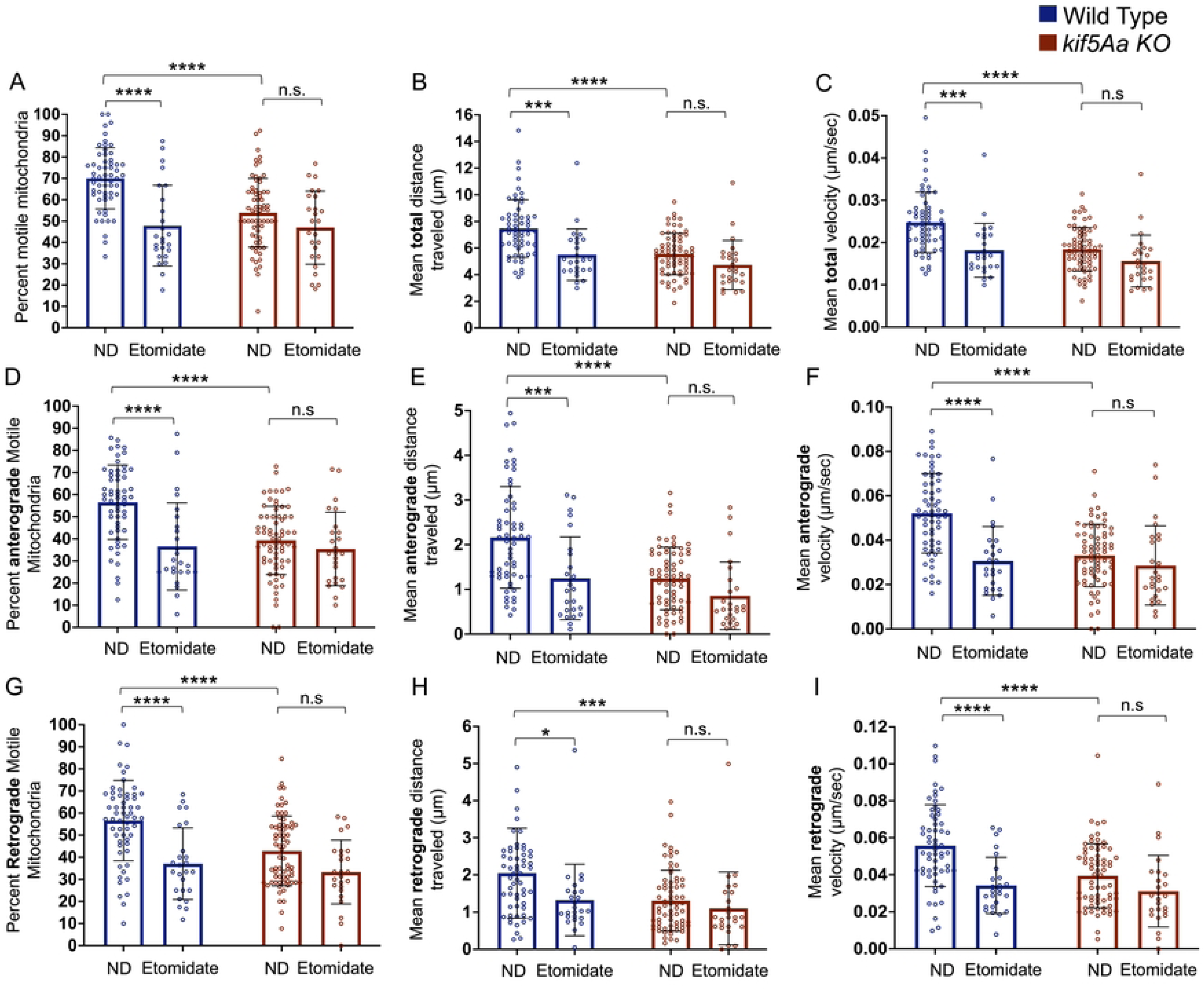
Exposure to etomidate decreases anterograde and retrograde mitochondrial motility in WT neurons. Total motility (A-C) was assessed and then broken into anterograde (D-F) and retrograde (G-I) movement. A) There was a significant decrease in motile mitochondria within the WT cells (blue bars) as well as a decrease in total distance moved (B, blue bars) and velocity (C, blue bars). When assessing anterograde movement, there was a decrease in the number of motile mitochondria (D, blue bars), the distance moved (E, blue bars), and the velocity (F, blue bars). Additionally, when assessing retrograde motility, we found a decrease in the number of motile mitochondria (G, blue bars), distance moved (H, blue bars), and velocity (I, blue bars). A-I) However, within the *kif5Aa* KO neurons, there was no significant change in motility when exposed to etomidate (red bars). n= 25-68 (see Supplemental Table 2). Error was in standard deviation and statistical significance using a two-way ANOVA with multiple comparisons using Bonferroni’s correction.

In contrast, dexmedetomidine produced no significant changes in mitochondrial motility or velocity from WT neurons (**Fig. 7A-I**, blue bars). However, a trend existed for increased distance moved and velocity in the retrograde direction (**Fig. 7H-I**, blue bars). If we focused on the motile mitochondria (56% of total), dexmedetomidine increased the distance moved and velocity (**Fig. 7J-K**).

**Figure 7.**
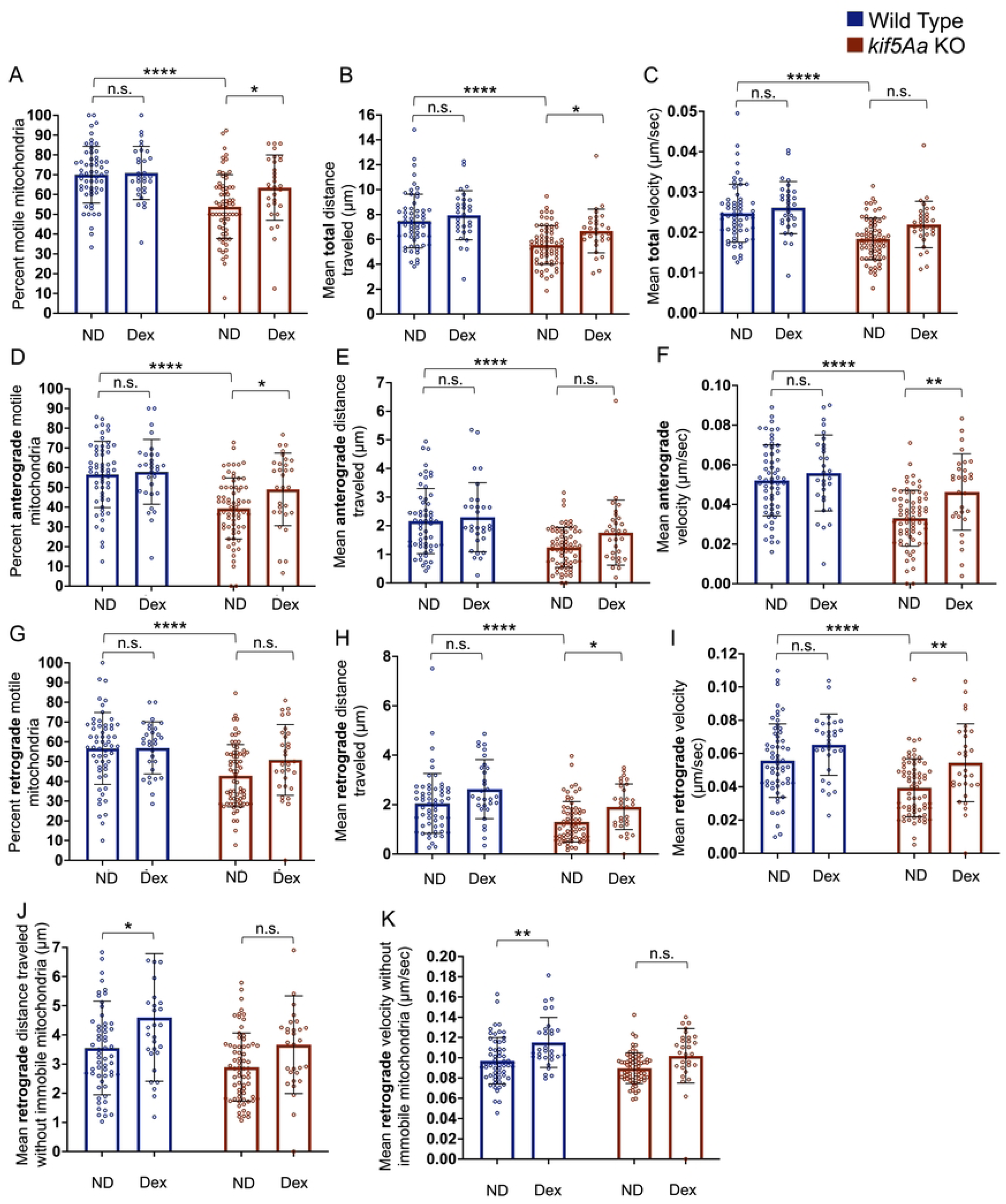
Exposure to dexmedetomidine increased mitochondrial motility in *kif5Aa* KO neurons. A-I) There was no significant change in mitochondrial motility (blue bars) when WT neurons were assessed. There was a significant increase in motile mitochondria within the *kif5Aa* KO cells (A, red bars) and in the total distance moved (B, red bars). However, the velocity (C, red bars) was not significantly different. When assessing anterograde motility, there was an increase in the number of motile mitochondria (D, red bars) and their velocity (F, red bars) but not in the distance they moved (E, red bars). Retrograde mitochondrial motility showed a similar number of motile mitochondria (G, red bars) but an increase in the distance moved (H, red bars) and velocity (I, red bars). When the immobile mitochondria were removed for the average, we assessed WT neuron mitochondria motility. We found a significant increase in the WT mitochondrial retrograde distance traveled (J, blue bars) and velocity (K, blue bars) but not *kif5Aa* KO mitochondria (J-K, red bars). n= 30-68 (see Supplemental Table 2). Error was in standard deviation and statistical significance using a two-way ANOVA with multiple comparisons using Bonferroni’s correction.

Within the *kif5Aa* KO neurons, dexmedetomidine exposure produced a significant increase in the total number of motile mitochondria (**Fig. 7A**, red bars), and distance moved (**Fig. 7B**, red bars), without changing velocity (**Fig. 7C**, red bars). Dividing motility into anterograde and retrograde in the *kif5Aa* KO neurons, dexmedetomidine caused significantly more mitochondria to move anterograde (**Fig. 7D**, red bars), coupled with an increase in velocity (**Fig. 7F**, red bars), but without a significant change in the distance traveled (**Fig. 7E**, red bars). Dexmedetomidine also caused a significant increase in both the retrograde distance moved (**Fig. 7H**, red bars) and the velocity (**Fig. 7I**, red bars) of the mitochondria. When focusing on the motile mitochondria, we saw no significant increase in either distance traveled (**Fig. 7J**, red bars) or velocity (**Fig. 7K**, red bars).

Similar to dexmedetomidine, ketamine produced no significant alteration in the motility of WT neuronal mitochondria (**Fig. 8A-I**, blue bars), although there was an increase in the number of motile mitochondria within the *kif5Aa* KO neurons (**Fig. 8A**). However, this did not lead to any change in distance moved or velocity (**Fig. 8B-C**). In the *kif5Aa* KO neurons, ketamine increased anterograde mitochondrial motility and velocity (**Fig. 8D, F**) but no change was detected in the distance moved (**Fig. 8E**). Finally, ketamine produced no change in retrograde mitochondrial motility in the *kif5Aa* KO neurons (**Fig. 8G-I**).

**Figure 8.**
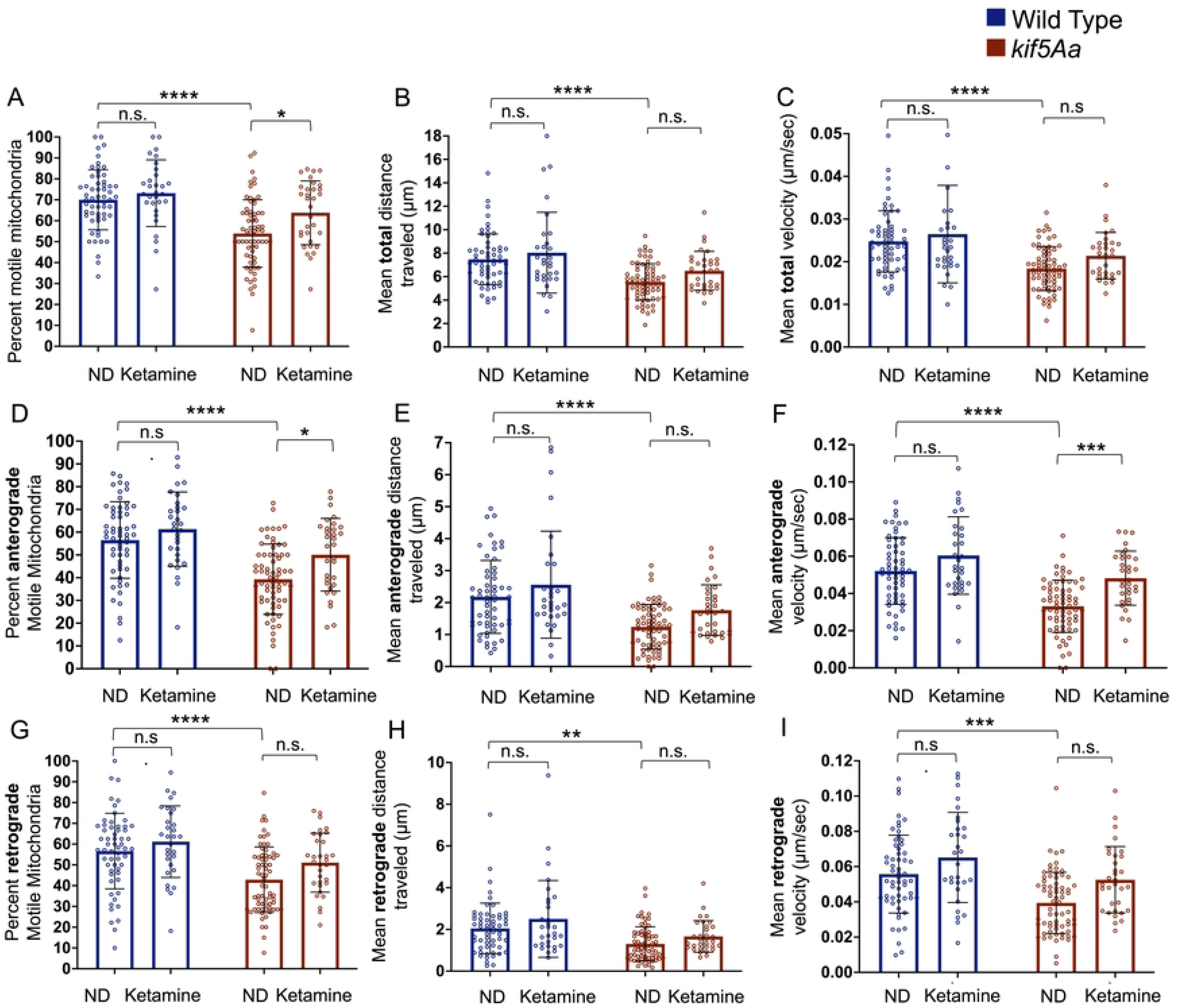
Exposure to ketamine causes minimal changes in mitochondrial motility. A-I) Within WT neurons, there was no significant change in mitochondrial motility in response to ketamine (blue bars). Within the *kif5Aa* KO neurons, there was an increase in the total number of motile mitochondria (A, red bars), with that increase being primarily anterograde movement (D, red bars) and not retrograde (G, red bars). B-D, red bars) There was no change in the total distance moved or velocity. There was an increase in anterograde velocity (F, red bars) but no change in the distance moved (E, red bars). G-I) There was no change in retrograde mitochondrial movement within the *kif5Aa* neurons. n= 30-68 (see Supplemental Table 2). Error was in standard deviation and statistical significance using a two-way ANOVA with multiple comparisons using Bonferroni’s correction.

## Discussion

Here we use the zebrafish model organism to expand upon the effect various IV anaesthetic/sedative drugs have upon intracellular motors, specifically Kif5A, as reflected by mitochondrial motility. When comparing human Kif5B with zebrafish Kif5Aa, there is a 67% overall homology, with very high homology in the motor domain (**Supplemental Fig. 1**, blue line). Additionally, residues defining the previously published propofol binding site[26] are conserved (**Supplemental Fig. 1**, red boxes). However, previous studies were performed *in silico*, *in vitro*, or in cell culture[15–17, 26]. Therefore, it was unknown if propofol binding to Kif5A was associated with altered behavioural responses to sedatives/anaesthetics. Additionally, since a known cargo of Kif5A is mitochondria, it was unclear whether IV drugs altered their distribution in the neuron, with potentially important energetic consequences.

Here, we provide data suggesting that mitochondrial transport is important in the behavioral sensitivity to certain sedation/anesthetic drugs. Our behavioral data showed that the *kif5Aa* KO larvae were significantly more sensitive to the GABAeric drugs, propofol, and etomidate compared to their WT siblings and that sensitivity to etomidate was greater than to propofol. This sensitivity within the mutant animals was consistent with the *in vitro* data. It had been shown that, for propofol, the motor domain of kinesin was bound, which induced the motor protein to fall off the microtubule. This, theoretically, caused a decrease in energy available at the synapse. However, within the *kif5Aa* KO larvae, there was already a dearth of mitochondria at the synapse. Therefore, the effect of this binding was accounted for prior to induction. Thus, the energy-poor synapses would be more vulnerable to the other mechanisms of the anesthetics.

Unexpectedly, the KO animals’ sensitivity to the other drugs, ketamine and dexmedetomidine, was less evident; in fact, we noted a decrease in sensitivity to the alpha 2 agonist dexmedetomidine. The complete lack of a behavioral phenotype to ketamine was unanticipated. However, it is still unknown if the kinesin motor domain is bound and inhibited by ketamine. Alternatively, the *kif5Aa* KO larvae were resistant to dexmedetomidine. This suggests other protein binding partners within the motor protein pathway for these compounds.

When considering neuron energy homeostasis, previous studies within mice on the loss of a critical protein in the electron transport complex 1 (ndufs4), caused an increase in sensitivity to both propofol and a resistance to ketamine[27]. Therefore, if the *kif5Aa* motor protein phenotype was only due to energetics, this data would be inconsistent. However, unlike the ndufs4 protein, altering motor protein transport also altered mitochondrial health and influenced fission/fusion events within the neuron. Adding to the complexity, mitochondria are not the only anaesthetic-bound protein/organelle to be transported by the kif5A protein family. In fact, it has been shown that kif5A is critical in the transport of the GABAA receptor[28]. Therefore, the alteration of neuronal subcellular processes was likely complex and compound-specific.

Our *in vivo* studies used two fundamentally different behavioral endpoints to study anesthetic/sedative compounds. These two endpoints responded differently over time. Within the GABAergic drugs, propofol and etomidate, SM was consistent, whereas EM had a time-dependent phenotype. SM was a more complex behavior that involved decision-making circuits within the larvae[29]. However, given the complexity, it would theoretically be easier for a drug to disrupt those neural circuits and thus may explain why we see a consistent phenotype over time. Alternatively, the neural circuits EM has been elucidated. This behavior was a less complex but evolutionarily critical reflex arc[30–32]. Theoretically, these critical reflex neurons would require more time to run out of energy. Therefore, we see a slow increase in sensitivity over time.

Consistent with the *in vitro* single kinesin motor protein and binding studies[15, 16], mitochondria within WT neurons moved less when exposed to both GABAergic drugs. Propofol produced anterograde movement inhibition, whereas etomidate inhibited both anterograde and retrograde movement. These results suggest an additional effect on dynein and/or its accessory proteins. Consistent with dexmedetomidine and ketamine acting on different molecular targets than etomidate or propofol, they had a very different effect on mitochondrial transport. Ketamine had little effect, while dexmedetomidine enhanced mitochondrial motility, especially in the retrograde direction.

Knocking out *kif5Aa* reduced mitochondrial anterograde transport, and the incremental effect of adding the GABAergic anesthetics was eliminated. This suggests that Kif5Aa may be the key anterograde neuronal motor protein targeted by those drugs. Despite the decrease in motility, anterograde transport is still seen within the KO cells. Therefore, other kinesin proteins may compensate for the loss of the single kinesin but incompletely. For example, while Kif5Aa appears to be the important motor protein for neuron mitochondria, there is a similar protein, Kif5Ab. Therefore, since in *kif5Aa* KO fish, *kif5Ab* may be upregulated. Alternatively, *kif1* and *klp6* families also contribute to neuron mitochondrial anterograde motility[12–14]. Nevertheless, despite neurons containing kinesins that may partially compensate for the loss of Kif5Aa, when the *kif5Aa* KO neurons were exposed to propofol and etomidate, there was no further decrease in mitochondrial motility. This suggests that neither propofol nor etomidate targets these alternative kinesins.

Retrograde motility is dependent upon dynein motor proteins[33, 34]. To our knowledge, the effect of anesthetics/sedatives on this key neuronal regulatory protein has not been studied. Here, we show that etomidate and dexmedetomidine influence retrograde motility. With etomidate, there is a decrease in WT mitochondria retrograde motility. Additionally, while not significant, there is a trend toward a decrease in motility within the *kif5Aa* neurons that is not seen in propofol, suggesting that a subset of neurons may be affected by etomidate. Compared to etomidate, dexmedetomidine does not alter the number of motile retrograde mitochondria within WT cells but does increase their distance and speed. Additionally, there is an increase in mitochondrial retrograde motility within the *kif5Aa* KO cells. This suggests that dexmedetomidine enhances mitochondrial transport from the synapse to the soma. It is still unclear if this alters synaptic mitochondrial health.

When interpreting these data, there are two key considerations. First, these data were collected from neurons cultured from whole brains, which include many different neuronal populations: motor, sensory, inhibitory, and excitatory. Given this heterogeneity, it is unsurprising that we observe large variability when assessing neuron mitochondrial motility. It is still unknown if the mitochondrial motility phenomenon is pan-neural or if different neurons are differentially affected. Therefore, the next step is to fully utilize the power of the zebrafish model by obtaining animals with fluorescently labeled neuronal subsets and assessing motility within those various classes.

Second, it is important to consider that the nervous system of 5 dpf fish is still growing and making critical connections. Therefore, the brain may respond differently to changes in motor and energetic homeostasis compared to the adult brain. It is still unknown if the mechanisms described above will continue into adulthood.

In conclusion, anesthetic/sedative drugs influence whole animal behavior and mitochondrial motility similarly. Using a *kif5Aa* knock-out animal has enabled the conclusion that this class of motor protein produces changes in mitochondrial motility, and some anesthetic effects may be translated through these proteins. These data suggest the hypothesis that alteration of mitochondrial dynamics and distribution contribute to the anesthetic state.

## Details of authors’ contributions

PD: planned and conducted aspects of all the experiments and helped write and edit the manuscript. RD: analyzed some of the mitochondrial movement data. RGE: helped plan experiments and edited the manuscript. VMB: planned and conducted aspects of all experiments, analyzed the data, and wrote the manuscript.

## Acknowledgments

Dr. Florence Marlow (Icahn School of Medicine at Mount Sinai) kindly provided the *kif5Aa^sa7168^* KO line. Drs. Philip Morgan and Margaret Sedensky for their input on the manuscript.

## Declaration of generative AI and AI-assisted technologies in the writing process

During the preparation of this work, the author(s) used Grammarly to improve the document’s grammar. After using this tool/service, the author(s) reviewed and edited the content as needed and took (s) full responsibility for the content of the publication.

